# A clinically identified Kirrel3 variant disrupts trans-synaptic circuit assembly

**DOI:** 10.64898/2026.10.20.760423

**Authors:** Omar Shennib, Alyssa Johnson, Olivia Raines, Megan E. Williams

## Abstract

Trans-synaptic adhesion molecules guide selective connections between neurons to establish circuit specificity. Yet, there are gaps in the understanding of how many synaptic adhesion molecules function in vivo and if clinically identified variants perturb their function. Using cell-type-specific deletion in the mouse hippocampus, we show that Kirrel3 is required in both pre-synaptic DG neurons and post-synaptic GABA neurons to form DG-to-GABA excitatory synapses that drive a feed-forward inhibitory connection to CA3. These data provide in vivo, genetic evidence for a homophilic trans-synaptic Kirrel3 interaction driving synapse formation. Moderately increasing or decreasing wildtype Kirrel3 expression did not detectably alter synapse formation or CA3 neuron activity. In contrast, expression of a Kirrel3 missense variant identified in an individual with autism dominantly disrupts synapse formation and CA3 activity in vivo, providing a mechanism by which a single missense allele could disrupt circuit assembly and contribute to disease.

## INTRODUCTION

Cell adhesion molecules play essential roles in brain development, particularly in the formation, function, and plasticity of synapses^1–3^. The large number and diversity of cell adhesion molecules, along with their highly specific homophilic and heterophilic binding interactions, has led to the hypothesis that they could constitute a molecular “code” that guides synaptic specificity through lock-and-key logic^4–7^. Consistent with a role in the form and function of specific circuits, genetic variation in synaptic adhesion molecules is frequently associated with neurodevelopmental disorders and mental illnesses^1,8–12^. However, for most cell adhesion molecules, little is known about how changes in gene dosage or expression of clinically identified missense variants affect synapse formation in vivo. This limits the ability of clinicians to determine if a genetic variant could be causative of disease.

Kirrel3 is an excellent model for addressing these knowledge gaps. It is a single-pass transmembrane protein from the immunoglobulin superfamily that promotes synapse formation across species^13–19^. Mammalian Kirrel3 protein is homophilic and binds itself in trans^20–23^. In the mouse hippocampus, Kirrel3 is selectively expressed by dentate gyrus (DG) granule neurons and a subset of GABAergic (GABA) neurons, where it is required for the formation of a specific type of DG synapse^22,24,25^. DG axons form large mossy fiber boutons that synapse onto CA3 pyramidal neurons. In addition, filopodia extend from these boutons to synapse onto nearby GABA neurons, which in turn provide feed-forward inhibition to CA3 (Fig. 1A,B)^26–28^. Kirrel3 is specifically required for these DG-to-GABA filopodial synapses. Germline knockout mice lacking Kirrel3 from all cells exhibit a marked reduction in these connections and increased CA3 activity, consistent with impaired feed-forward inhibition^20,24^.

**Figure 1.**
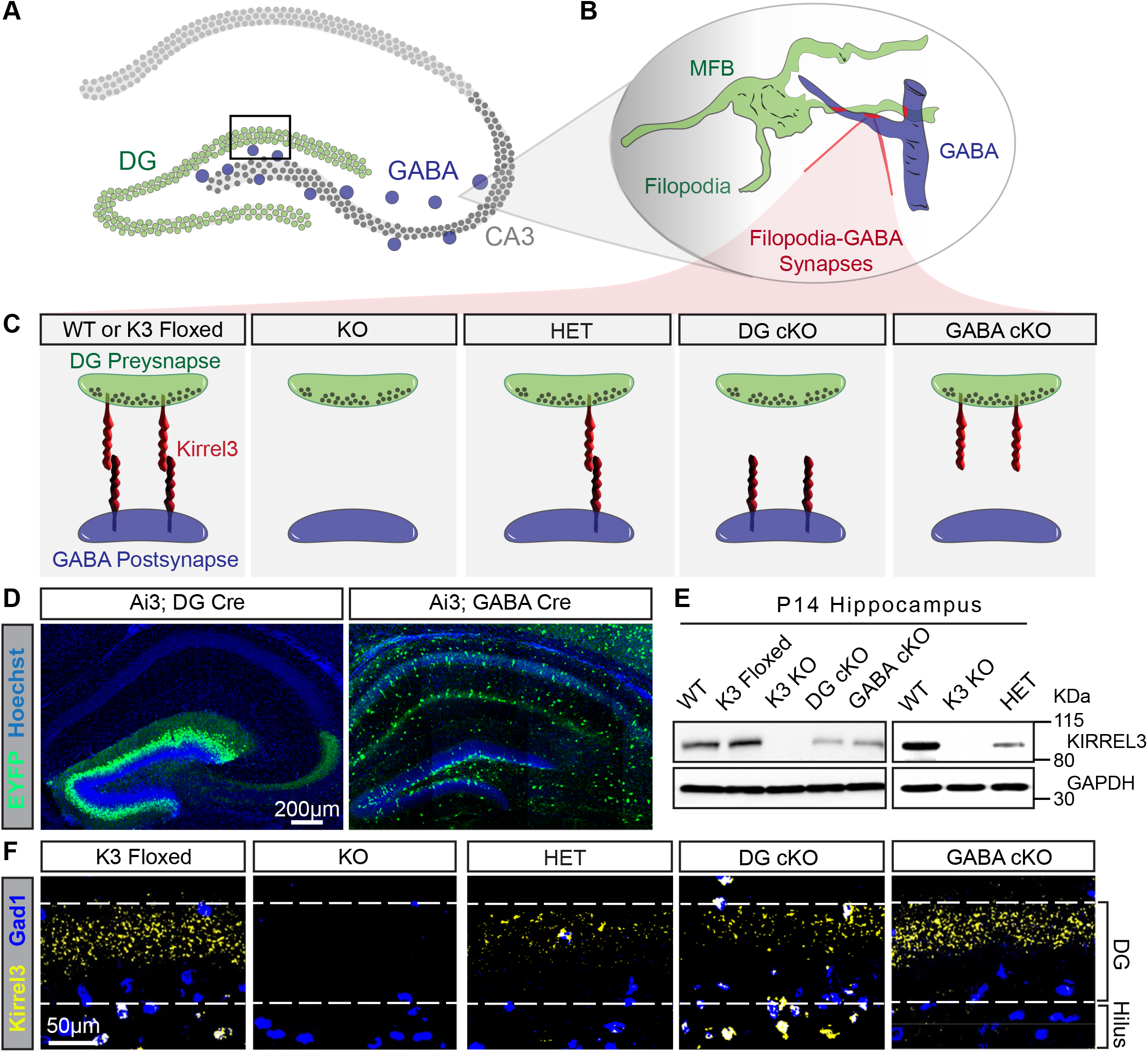
Cell-type-specific deletion of Kirrel3 from DG or GABA neurons. **A)** Hippocampus diagram. The box marks the region of interest shown in F. **B)** Mossy fiber bouton and filopodia-GABA synapse diagram. **C)** Diagram of differential Kirrel3 protein in transgenic mice. **D)** DG Cre and GABA Cre mice crossed to the Ai3 EYFP Cre-reporter line indicate that Cre is expressed in the intended cell types. Hippocampal sections from P14 mice. Images are tiled. **E)** Western blots of P14 hippocampal lysates with indicated antibodies. **F)** Examples from the DG soma layer of sections used for fluorescence in situ hybridization for mRNAs of Kirrel3 (yellow), and the GABA neuron marker Gad1 (blue). Dash lines indicate DG cell body layer as determined by Hoechst (not shown). Note that in the DG area of K3^flox/flox^ (K3 floxed) mice, Kirrel3 is normally expressed in the outer DG layer where postmitotic DG granule neurons reside, and it is not expressed in the subventricular zone and lower layers containing stem cells and immature neurons. Kirrel3-expressing GABA neurons are shown here in the DG region but they are also found in all hippocampal regions.

In vitro, Kirrel3 is sufficient to induce synapses when ectopically expressed^22^, but whether Kirrel3 is required in both pre- and postsynaptic neurons for synapse formation in vivo has not been tested. In addition, human genetic studies identified numerous missense variants and copy number variations in Kirrel3 in individuals with neurodevelopmental disorders and mental illnesses^10,22,29–34^. Most individuals with Kirrel3 variants are heterozygous, but the functional consequences of these variants, and the sensitivity of synapse formation to Kirrel3 heterozygosity or gene dosage are unknown. Here, we define the cell type–specific and gene dose– dependent requirements for mouse Kirrel3 and two clinically-identified variants in hippocampal synapse formation using conditional genetic and viral approaches. We find that Kirrel3 must be expressed by the pre and postsynaptic neuron, supporting a trans-cellular role in vivo. We also find synapse formation tolerates modest increases and decreases in Kirrel3 expression, but synapse formation in vivo is strikingly sensitive to expression of a rare missense variant identified in an individual with autism spectrum disorder^33^. This work identifies a mechanism by which a single mutant Kirrel3 allele can impair circuit assembly.

## RESULTS

### Validation of Kirrel3 conditional knockout mouse models

In the mouse hippocampus, Kirrel3 is expressed by DG neurons and some GABA neurons (Fig. 1A) and is required for DG-to-GABA synapses (Fig.1B)^20,24^. Thus, to define the cell type requirements of Kirrel3 in hippocampal synapse formation in vivo, we sought to generate and validate a set of conditional knockout (cKO) mice. We crossed a Kirrel3 floxed mouse line^35^ with Dock10-Cre line^36^ to generate a DG neuron specific cKO. We crossed the Kirrel3 floxed line with a Gad2-IRES-Cre line^37^ to generate a GABA neuron specific cKO. We also obtained germline Kirrel3 wildtype, heterozygous, and knockout mice^38^ (Fig. 1C). We first confirmed the cell type specificity of each Cre driver by crossing them to a Cre-dependent EYFP reporter line^39^. As expected, Dock10-Cre drives recombination specifically in mature DG neurons (Fig. 1D). Gad2-Cre labels scattered neurons throughout the hippocampus (Fig. 1D), consistent with the expected location of GABA neurons. We next assessed Kirrel3 protein levels in the hippocampus of each line at postnatal day 14 (P14), a time point when germline Kirrel3 knockout mice show a synaptic phenotype^20^. Western blot reveals a complete loss of Kirrel3 in germline knockout mice and a reduction to approximately 50% in heterozygous (Kirrel3^+/-^), DG cKO (Kirrel3^flox/flox^; Dock10^Cre/+^), and GABA cKO (Kirrel3^flox/flox^; Gad2^Cre/+^) animals (Fig. 1E). To validate the cellular specificity of Kirrel3 deletion in the cKOs, we conducted fluorescent in situ hybridization at P14. In wildtype and Kirrel3 floxed mice without a Cre driver, Kirrel3 mRNA is detected in DG neurons and in Gad1-positive GABA neurons (Fig. 1F and Supplement to Fig. 1). Kirrel3 transcripts are absent in germline knockout mice and heterozygotes show reduced Kirrel3 mRNA signal from both DG and GABA neurons (Fig. 1F). DG cKO mice have a marked, but incomplete removal of Kirrel3 mRNA in the DG, with intact expression in GABA neurons (Fig. 1F). Conversely, GABA cKO mice show a loss of Kirrel3 mRNA from GABA neurons, with intact DG expression (Fig. 1F). Together, these results validate a set of Kirrel3 mouse models that enable cell type specific and gene dose dependent interrogation of Kirrel3 function in synapse formation and circuit activity.

### Mossy fiber filopodia synapse formation requires Kirrel3 expression in both pre-synaptic DG and postsynaptic GABA neurons

To determine if Kirrel3 is required in presynaptic DG neurons, postsynaptic GABA neurons, or both, we quantified DG-to-GABA mossy fiber filopodia synapses in DG and GABA cKO mice at P14. To visualize and quantify individual DG mossy fiber filopodial synapses, we labeled DG mossy fiber axons by injecting a membrane-targeted EGFP AAV into neonatal pups. Sparse labeling enabled high-resolution three-dimensional visualization of individual axons, boutons, and their filopodia (Fig. 2A). Synapses were identified by immunostaining for synaptoporin, a synaptic vesicle marker enriched at DG presynapses^40,41^ (Fig. 2A). Discrete synaptoporin puncta within EGFP-labeled filopodia were quantified per mossy fiber synapse complex as a measure of synapse density (Fig. 2A–C). Both DG and GABA cKO mice exhibited a significant reduction in mossy fiber filopodial synapses compared to Kirrel3 floxed only controls (Fig. 2B-D). DG cKO mice showed an approximately 50% decrease in filopodial synapses per bouton, whereas GABA cKO mice exhibited a pronounced ∼75% reduction (Fig. 2D). Notably, this large reduction in GABA cKOs is very similar to prior ultrastructural analyses reporting ∼80% fewer filopodia synapses per bouton in Kirrel3 germline knockout mice^24^. Thus, loss of Kirrel3 from only GABA neurons phenocopies the germline knockout in the hippocampus. The less severe reduction in DG cKO mice is consistent with incomplete recombination and residual Kirrel3 expression in this Cre line at this age (Fig. 1F). These results demonstrate that Kirrel3 must be expressed in both presynaptic DG neurons and postsynaptic GABA neurons for normal formation of mossy fiber filopodial synapses. This supports a model in which Kirrel3 uses trans-synaptic, homophilic interactions to form synapses selectively between Kirrel3-expressing neurons in the brain.

**Figure 2.**
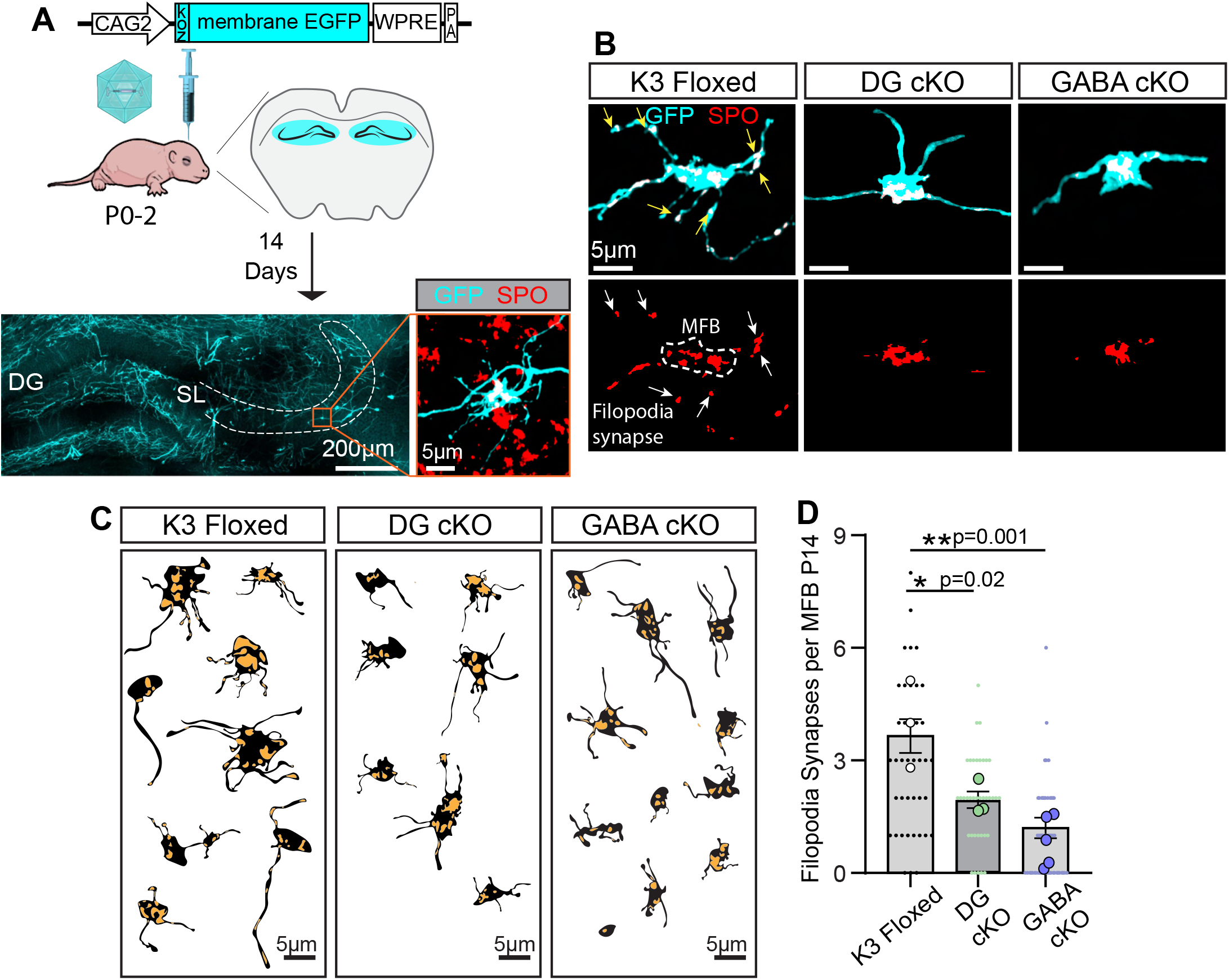
Kirrel3 is required in DG and GABA neurons for mossy fiber filopodia synapse formation. **A)** Diagram of the mossy fiber synapse labeling assay using the indicated AAV with an example of labeled DG neurons and a magnified image of a membrane-EGFP labeled mossy fiber synapse (cyan) stained with a synaptoporin antibody (SPO, red). SL; stratum lucidum mossy fiber layer. **B)** Confocal images of a labeled mossy fiber bouton (MFB), its filopodia, and SPO immunostaining for indicated genotypes. Note that endogenous SPO staining outside of the GFP signal was removed for visualization. **C)** Tracings of representative mossy fiber synapses in black with associated SPO synapses in yellow. **D)** Quantification of the average number of filopodia synapses (SPO puncta) per MFB. Sample sizes as follows: n= 43 MFBs from 3 mice (Floxed), 37 MFBs from 3 mice (DG cKO), and 72 MFBs from 5 mice (GABA cKO). Nested one-way ANOVA followed by pairwise post-tests. Large dots indicate the average filopodia synapse per mouse. All mice were P14 including both sexes.

### Kirrel3 expression is required in DG and GABA neurons to maintain normal CA3 activity patterns

Mossy fiber filopodial synapses provide feed-forward inhibition to CA3 neurons, and their loss is associated with increased CA3 activity^20,26,27,42^. To test if the decrease in filopodia synapse density observed in DG and GABA Kirrel3 cKO mice alters circuit activity, we measured overall CA3 activity in each genotype using cFos immunohistochemistry^43^. At P14, all genotypes tested have a similarly low density of cFos-positive CA3 neurons when kept in their home cage (Supplementary Fig. 3A), indicating no overt differences in resting activity. To broadly stimulate the hippocampus, we subjected mice to a mild foot shock^28,44,45^, and then quantified cFos expression (Fig. 3A). After foot shock, GABA cKO mice have a significant increase in the density of cFos-positive CA3 neurons compared to controls (Fig. 3B,C). This robust increase in active CA3 neurons in GABA cKO mice is similar to prior observations in germline Kirrel3 knockout mice^20^. The increased CA3 activity in GABA cKO mice, together with the loss of filopodial synapses in GABA cKO mice (Fig. 2D), indicates a reduction of feed-forward inhibition to CA3. By contrast, DG cKO mice exhibited normal CA3 activity at this age (Fig. 3B,C). Normal CA3 activity in DG cKO mice is consistent with their more modest reduction in filopodial synapse density (Fig. 2D) and indicates that the remaining synapses in DG cKO mice are sufficient to maintain CA3 activity at the population level as measured by cFos.

**Figure 3.**
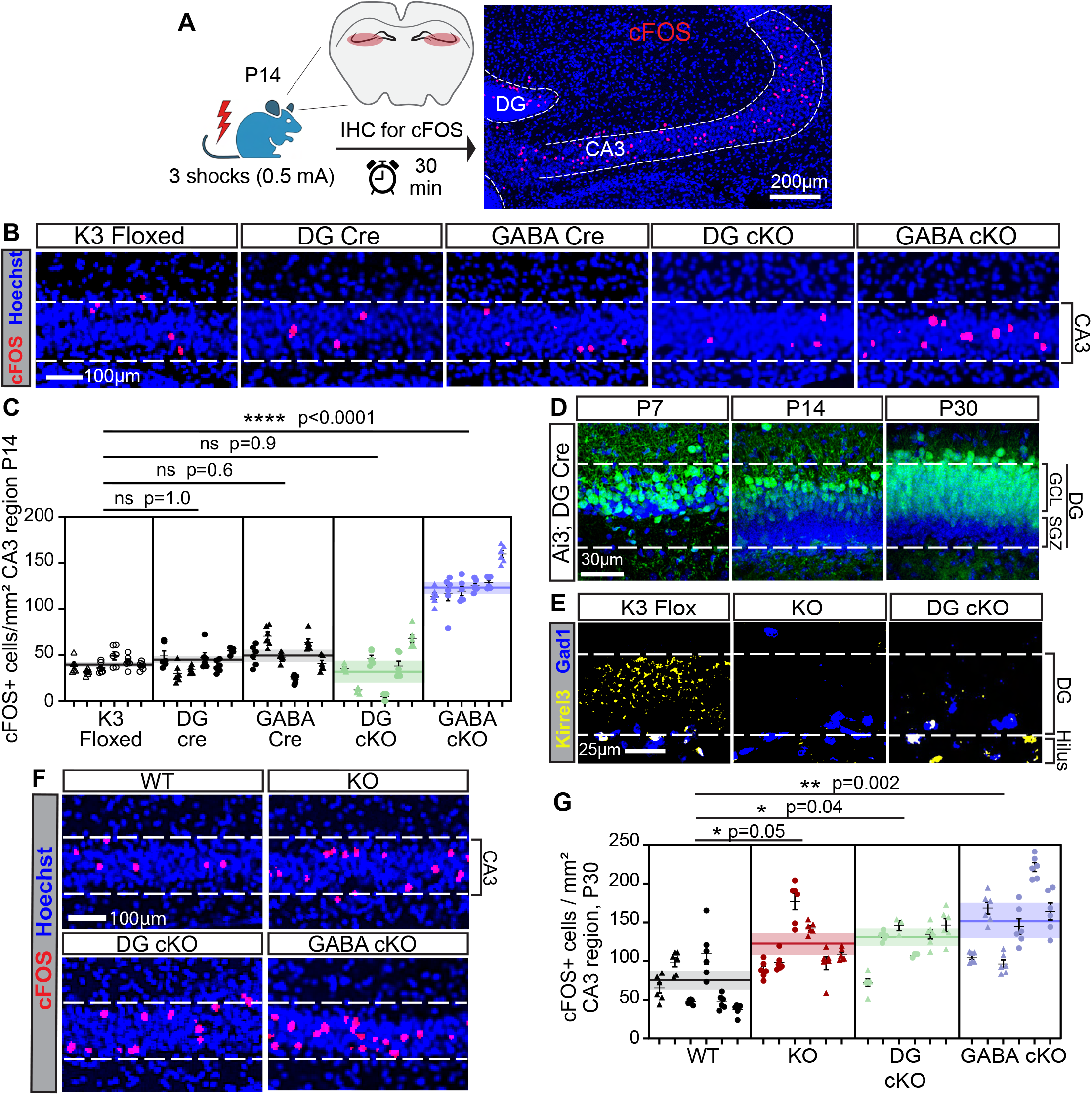
Kirrel3 is required in DG and GABA neurons to maintain normal CA3 activity. **A)** Diagram of the foot shock assay and example cFOS immunostaining in CA3 pyramidal layer (dashed lines). cFOS (red), Hoechst (blue). **B)** Representative images of cFOS-positive (cFOS+) CA3 cells for each genotype after foot shock at P14. **C)** Quantification of the density of cFOS+ CA3 neurons after foot shock. n = 6 sections per mouse and 6 mice per genotype with males a triangle, females a circle. Data from each mouse is shown in a column. The horizontal line and shading across each genotype is the global average and SEM. Nested one-way ANOVA with pairwise post-tests as indicated. **D)** Images from the DG layer of Dock10-Cre (aka DG Cre) mice crossed to an Ai3/EYFP Cre reporter at indicated ages. EYFP positive neurons (green), Hoechst (blue), GCL, granule cell layer; SGZ, subgranular zone. **E)** Fluorescence in situ hybridization for mRNAs of Kirrel3 (yellow) and Gad1 (blue) in P30 mice. Dashed lines indicate DG cell body layer. **F)** Representative images of cFOS+ CA3 neurons for each genotype after foot shock at P30. **G)** Quantification of the density of cFOS+ CA3 neurons at P30. n = 3-6 brain sections per mouse and 6 mice per genotype with males represented by triangle and females a circle. The horizontal line and shading across each genotype is the global average and SEM. Nested one-way ANOVA with pairwise post-tests as indicated.

We initially examined cKO mice at P14 because our prior work on germline KO mice showed a clear phenotype at this age. However, because Kirrel3 mRNA was incompletely reduced by the Dock10-Cre line at P14 (Fig. 1F), we determined the developmental time course of Dock10-Cre–mediated recombination using the Cre-dependent EYFP reporter. We find that Dock10-Cre recombination in DG neurons increases substantially by P30 (Fig. 3D). Consistent with this, fluorescent in situ hybridization at P30 reveals a near-complete loss of Kirrel3 mRNA in DG neurons with normal expression in GABA neurons (Fig. 3E). We therefore examined CA3 activity after foot shock at P30. At this age, germline KO mice have a noticeable increase in cFos-positive CA3 neurons in the unstimulated home cage condition (Supplemental figure 3B), but the difference is not statistically significant. Importantly, after foot shock, germline knockout mice, DG cKO, and GABA cKO mice all have an elevated density of cFos-positive CA3 neurons compared to control mice (Fig. 3F,G). Thus, once Dock10-Cre-mediated deletion was near complete, loss of Kirrel3 from either synaptic partner increased CA3 activity, consistent with reduced feed-forward inhibition of CA3 neurons.

### Kirrel3 heterozygosity does not detectably alter synapse formation or CA3 activity

Individuals with *KIRREL3*-associated neurodevelopmental disorders often have a copy number or missense variant in only one Kirrel3 allele. We therefore tested if Kirrel3 is haploinsufficient for hippocampal synapse formation in mice. To test this, we analyzed filopodial synapse formation and CA3 activity in germline Kirrel3 heterozygous animals. As a positive control, we analyzed germline Kirrel3 KO mice, which have a severe reduction in filopodia synapse density (Fig 4A-C) and a corresponding increase in the density of active CA3 neurons after foot shock (Fig 4D,E)^20^. Kirrel3 heterozygous mice have a lower mean filopodial synapse density than wildtype mice, but this difference is not statistically significant (Fig. 4A–C). Moreover, CA3 activity following foot shock is comparable between Kirrel3 wildtype and heterozygous mice (Fig. 4D,E). Thus, within the resolution of these assays, heterozygous loss of wildtype Kirrel3 does not detectably alter mossy fiber filopodial synapse density or CA3 activity at P14.

**Figure 4.**
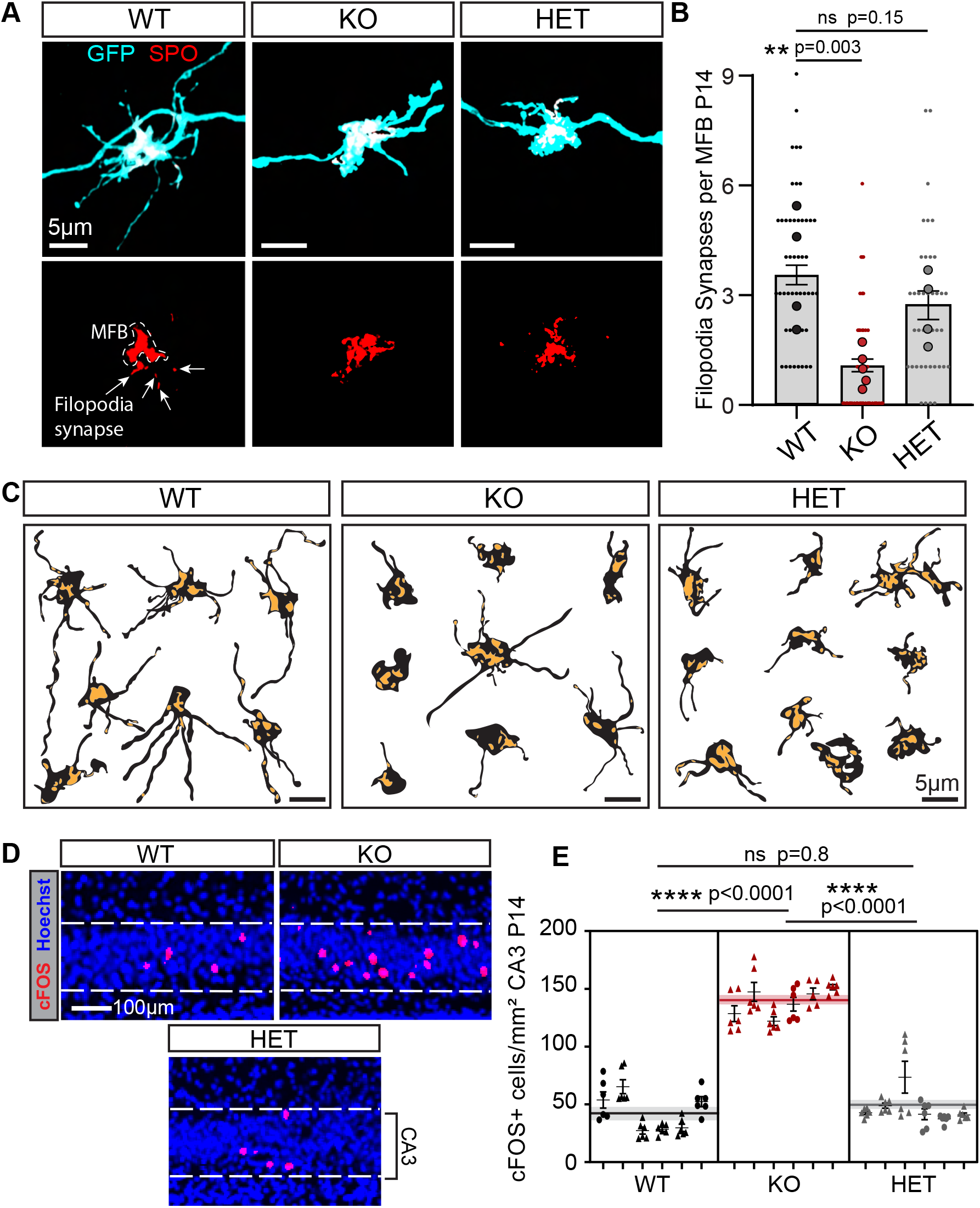
Heterozygous Kirrel3 loss does not detectably alter synapse density or CA3 activity. **A)** Representative images of MFBs and filopodia. EGFP (cyan) and SPO (red). The endogenous SPO from uninfected axons is masked. SPO puncta in filopodia are indicated by arrows. **B)** Quantification of the average number of filopodia synapses (SPO puncta) per MFB at P14. Sample sizes as follows: n= 49 MFBs from 4 mice (WT), 67 MFBs from 5 mice (KO), and 40 MFBs from 4 mice (HET). Nested one-way ANOVA followed by pairwise post-tests. Large dots indicate average filopodia synapse density per MFB per mouse. **C)** Tracings of representative MFBs with their SPO synapses shown in yellow. **D)** Representative images of cFOS+ CA3 cells in each genotype after foot shock at P14. **E)** Quantification of cFOS+ CA3 neurons at P14. n = 5-6 brain sections were quantified per mouse and 6 mice per genotype. Males are represented by a triangle and females a circle. The horizontal line and shading across each genotype is the global average and SEM. Nested one-way ANOVA with pairwise post-tests as indicated.

### A Kirrel3 variant dominantly impairs synapse formation and circuit activity

The dosage tolerance for heterozygosity in synapse formation drove us to test if clinically identified Kirrel3 missense variants act as dominant negatives in the presence of endogenous wildtype Kirrel3. Prior work showed that several clinically identified Kirrel3 missense variants have impaired synaptogenic activity in cultured neurons^22^. However, because the variants were tested in CA1 neurons that lack endogenous Kirrel3, those experiments could not determine if the variants are loss-of-function or dominant-negative alleles. We therefore tested their function in vivo in the presence of endogenous Kirrel3. We expressed wildtype Kirrel3 or missense variants in DG neurons of wildtype mice and assessed mossy fiber filopodia synapse density and CA3 circuit activity at P14. We selected two variants that were unable to induce synapses in culture: R161H, which retains a normal ability to mediate homophilic cell aggregation, and R205Q, which does not^22^. We confirmed that both mutants properly traffic to the cell surface of axons along with wildtype Kirrel3 (Fig. 5A). Next, we infected neonatal mice with AAVs expressing membrane-targeted EGFP to trace DG axons and filopodia, either alone as a control or together with HA-tagged Kirrel3 variants (Fig. 5B). We confirmed the AAVs induced expression of membrane GFP, and each Kirrel3 variant as expected (Fig. 5C, Supplement Fig. 5A-B). We then analyzed DG mossy fiber filopodia synapse formation using synaptoporin and CA3 activity after foot shock using cFos at P14.

**Figure 5.**
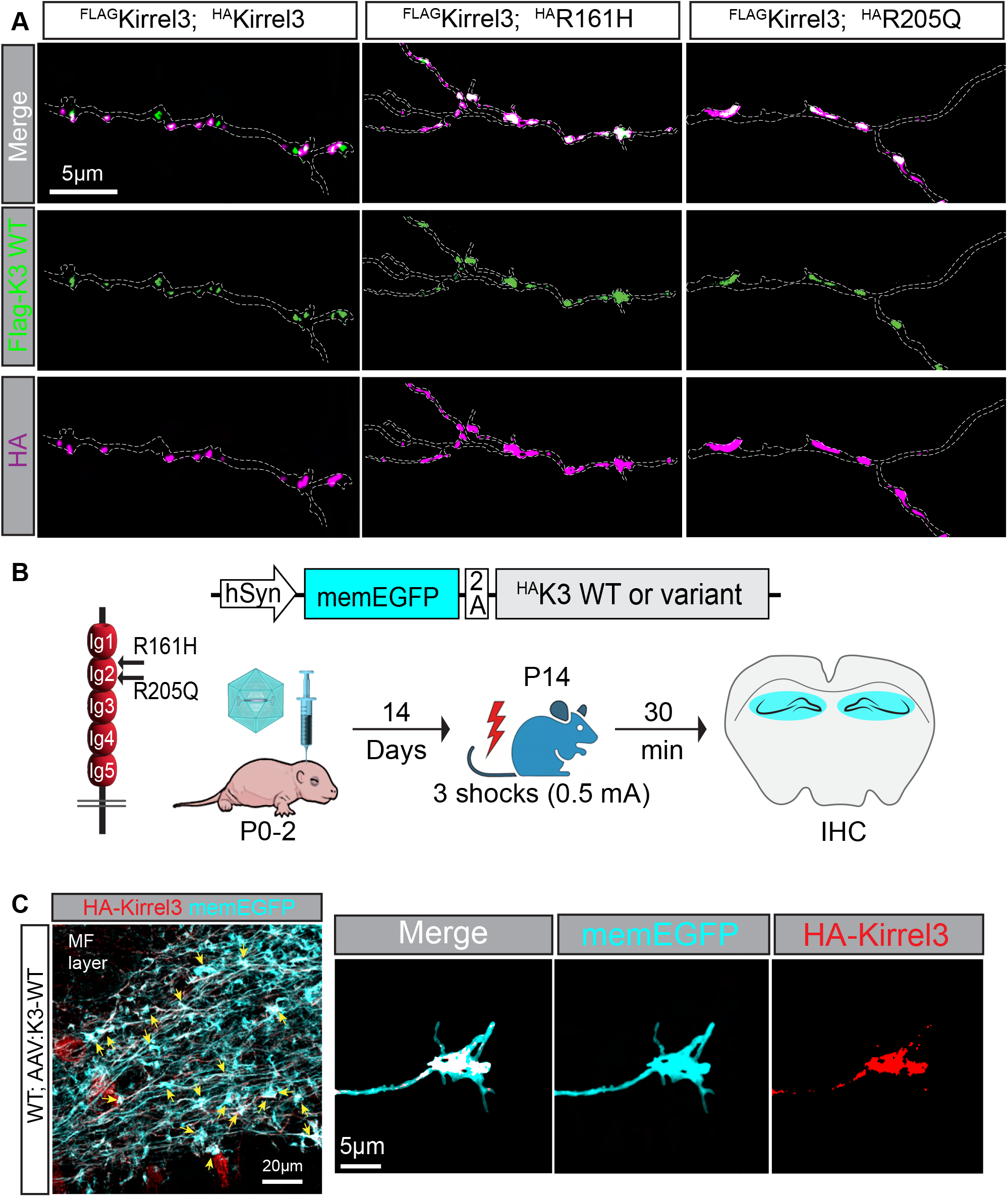
Kirrel3 variants show normal surface localization in axons. **A)** Representative images showing the colocalization of Kirrel3 wildtype and variants in cultured neuron axons. Hippocampal neurons were co-transfected with Flag (green) and HA (magenta) tagged Kirrel3 plasmids and live labeled to detect surface proteins. GFP (not shown) was also transfected and used to trace the axon indicated by dotted lines. **B)** Diagram of the variant location on Kirrel3 protein, AAV design, and overexpression workflow. After 14 days, alternating hippocampal sections were immunostained for cFOS, or GFP, HA-Kirrel3, and SPO. Ig, immunoglobulin domain. **C)** Example immunostaining showing expression of virally expressed membrane-EGFP (cyan) and HA-Kirrel3 (red) in DG mossy fiber axons.

Our results indicate that expression of wildtype Kirrel3 has no significant effect on filopodial synapse density or CA3 activity (Fig. 6A-E) compared to GFP controls. This indicates that increasing Kirrel3 expression in DG axons that normally express Kirrel3 does not adversely affect the circuit. Similarly, expression of the Kirrel3 variant R161H in wildtype mice had no significant effect on filopodia synapse density or CA3 activity. Because R161H was unable to induce extra synapses in cultured neurons that lack Kirrel3^22^, our results suggest that R161H is a loss-of-function or hypomorphic allele that does not interfere with wildtype Kirrel3 in vivo. In striking contrast, expression of the R205Q variant in wildtype mice markedly reduced filopodial synapse density and increased CA3 activity (Fig. 6A-E), with a magnitude resembling that of complete loss of Kirrel3 function. This indicates R205Q is a dominant-negative variant that interferes with endogenous Kirrel3 function.

**Figure 6.**
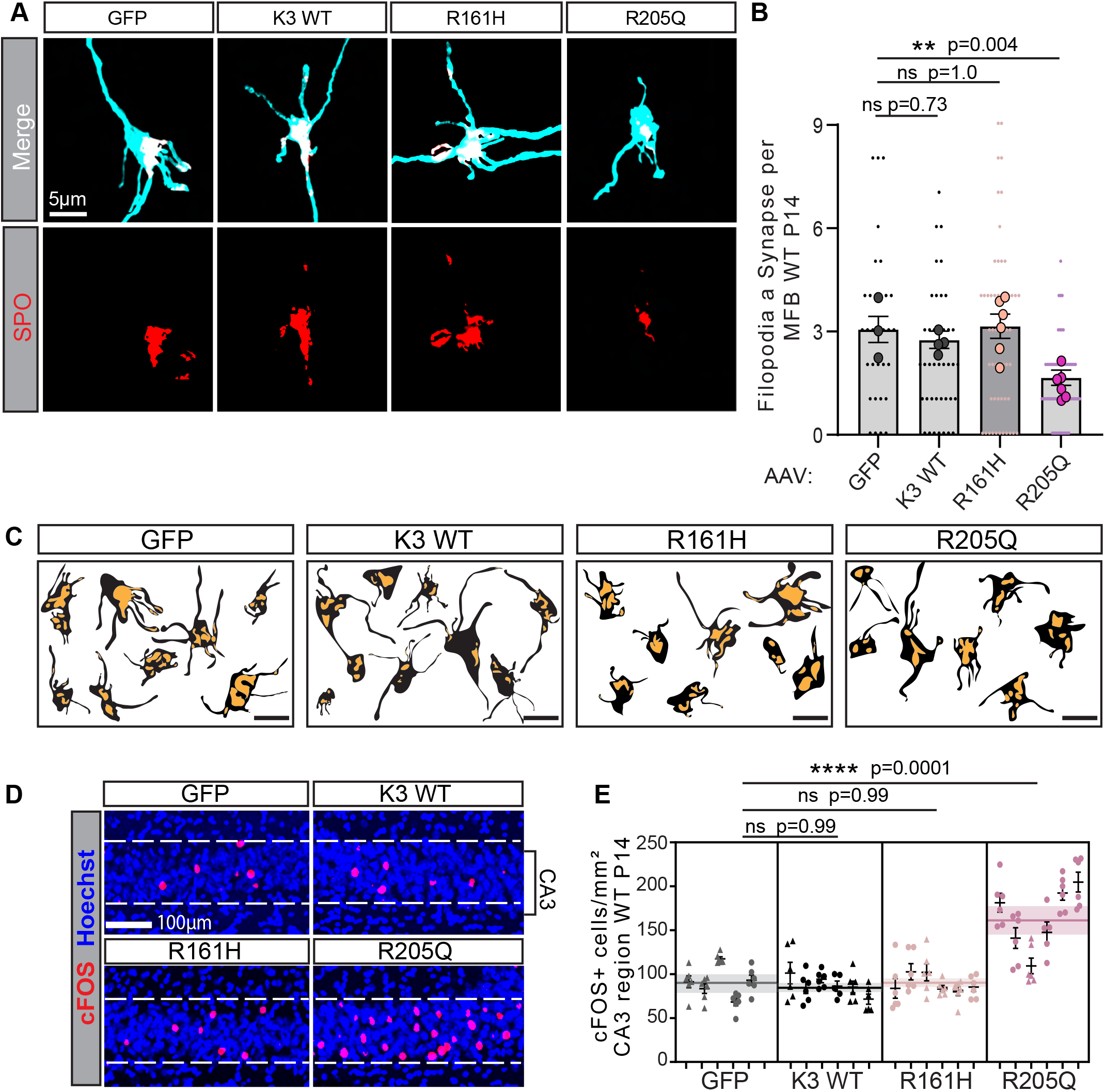
Expression of Kirrel3 R205Q impairs synapse formation and increases CA3 activity. **A)** Representative images of MFBs and filopodia. EGFP (cyan) and SPO (red). The endogenous SPO from uninfected axons is masked. **B)** Quantification of the average number of filopodia synapses (SPO puncta) per MFB at P14. Sample sizes as follows: n= 32 MFBs from 3 mice (GFP), 40 MFBs from 4 mice (K3 WT), 54 MFBs from 6 mice (R161H), and 62 MFBs from 6 mice (R205Q). Nested one-way ANOVA followed by pairwise post-tests. Large dot represents the average filopodia synapse density per mouse. **C)** Tracings of representative MFB and filopodia in black with SPO synapses in yellow. **D)** Representative images of cFOS+ CA3 cells in each genotype after foot shock. **E)** Quantification of the density of cFOS+ CA3 neurons after foot shock. n = 5-6 brain sections analyzed per mouse and 6 mice per genotype except GFP which has 5 mice. Males are indicated by a triangle and females a circle. The horizontal line and shading across each genotype is the global average and SEM. Nested one-way ANOVA with pairwise post-tests as indicated.

## DISCUSSION

Cell adhesion molecules are diverse and play essential roles in brain development, yet defining their precise functions in vivo has remained challenging^1,4^. This likely reflects the fact that their phenotypes are often subtle, cell type–specific, or context-dependent. Here, we address this challenge by defining how the homophilic adhesion molecule Kirrel3 regulates synapse formation in a defined hippocampal circuit and how genetic variation alters its function.

Our results demonstrate that mossy fiber filopodial synapse formation requires Kirrel3 expression in both presynaptic DG neurons and postsynaptic GABAergic neurons. Deletion of Kirrel3 from GABA neurons phenocopies the germline knockout, while partial deletion of Kirrel3 from DG neurons produces a partial phenotype. Loss of Kirrel3 from either synaptic partner also increases CA3 activity, linking disruption of these feed-forward inhibitory synapses to reduced inhibitory control of CA3 neurons. Interestingly, the DG cKO phenotype worsens with time as Dock10-Cre recombination increases. This suggests that even if mossy fiber synapse formation starts out normally, if Kirrel3 is removed from DG neurons in the first postnatal month, the density and function of the DG-to-GABA synapses is compromised. Together, our results provide in vivo evidence that Kirrel3 acts across synaptic partners to promote formation of a specific type of hippocampal synapse important for balancing activity in the hippocampus. These findings extend prior work demonstrating that Kirrel3 must be expressed in the pre and postsynaptic neuron for synapse formation in cultured neurons^22^ and that intact trans-cellular adhesion is necessary for proper formation of glomeruli in the vomeronasal organ^23^. Importantly, Kirrel3 is expressed by specific types of synaptically connected neurons throughout the brain^10,25,38^, raising the possibility that similar trans-synaptic interactions operate in other circuits. Thus, it is likely that Kirrel3 trans-cellular binding similarly mediates the formation of other types of synapses. Whether Kirrel3 acts during selective target selection, synapse formation, or stability remains to be determined in future work.

In contrast to complete Kirrel3 loss, heterozygous mice have a lower mean synapse density that is not statistically significant and no detectable change in CA3 activity. Thus, within the resolution of these assays, heterozygous Kirrel3 loss does not produce a detectable deficit in mossy fiber filopodial synapse formation or CA3 activity in mice. Similarly, viral-mediated overexpression of wildtype Kirrel3 did not alter synapse density or CA3 activity, indicating that this circuit tolerates moderate increases and decreases in wildtype Kirrel3 expression.

Many individuals with *KIRREL3*-associated disorders are not heterozygous loss of function but rather heterozygous for rare variants^10,22,29–32^. While prior in vitro assays identified variants with impaired synaptogenic activity, they could not distinguish between loss-of-function and dominant-negative variants^22^. By testing variants in the presence of endogenous wildtype Kirrel3, we identified a variant that dominantly disrupts synapse formation and circuit activity in vivo. R205Q, but not R161H, impaired synapse formation and increased CA3 activity despite wildtype Kirrel3 expression, supporting a dominant-negative effect of R205Q. Numerous rare Kirrel3 missense variants are reported in individuals with autism spectrum disorder, other neurodevelopmental disorders, and mental illnesses and different variants may affect circuit assembly through distinct cellular mechanisms. Testing additional variants will be needed to define the functional classes of other mutants. Nevertheless, our work suggests that variants that interfere with trans-binding (like R205Q) may act dominantly and our work establishes an in vivo framework for functionally classifying disease-associated Kirrel3 variants and copy number changes.

In sum, mossy-fiber filopodial synapse formation requires Kirrel3 in both synaptic partners, providing in vivo genetic evidence for a trans-synaptic homophilic mechanism. Although the circuit tolerated the tested changes in wildtype Kirrel3 dosage, the autism-associated R205Q variant disrupted synapse formation and increased CA3 activity despite endogenous wildtype Kirrel3. These findings provide a mechanistic link between a clinically identified missense variant and circuit dysfunction and suggest how a heterozygous KIRREL3 variant could contribute to neurodevelopmental disease.

## METHODS

### Mouse lines

All mice were maintained on a C57BL/6J background (RRID:IMSR_JAX:000664). All mouse lines were reported previously, including Kirrel3 floxed^35^, Dock10-Cre (RRID:MGI:6117432)^36^, Gad2-IRES-Cre (RRID:IMSR_JAX:010802)^37^, Kirrel3 knockout^38^, and the Ai3 Cre-dependent EYFP reporter (RRID:IMSR_JAX:007903)^39^. Mice were maintained according to NIH guidelines on the care and use of animals, and all procedures were approved by the University of Utah Institutional Animal Care and Use Committee. Details of key resources for all methods can be found in Supplementary Table 1.

### Immunohistochemistry

Mice were anesthetized with ketamine/xylazine (200 mg/kg:16 mg/kg) and transcardially perfused with 4% paraformaldehyde (PFA) in 1x phosphate-buffered saline (PBS) prepared from 10x stock consisting of 14.2 g of Na2HPO4, 80 g of NaCl, 2 g KCl, 2 g KH2PO4 at a pH of 7.4. Brains were then stored in 4% PFA overnight and sectioned at 50 μm through the dorsal hippocampus using a vibratome (Leica VT 1000S, RRID:SCR_016495). Free-floating sections were incubated in blocking solution (1x PBS with 3% bovine serum albumin (BSA), 0.2% Triton X-100 and 0.2% Saponin) for 1 hour. Sections were incubated overnight with gentle shaking in primary antibody solution, which was prepared by diluting primary antibodies in 1xPBS with 3% BSA. After 3 washes in 1x PBS, sections were incubated with secondary antibody solution for 2-3 hours at room temperature, followed by two washes in 1x PBS and a final wash in 1x PBS with Hoechst prior to mounting on slides in Fluoromount-G (Invitrogen Catalog# 50-187-88, RRID:SCR_015961).

### Antibodies

Primary antibodies were used as follows: goat anti-GFP 1:2000 (Abcam, ab6673, RRID:AB_305643), rabbit anti-GFP 1:2000 (Jackson ImmunoResearch Laboratories, 300-005-245, RRID:AB_3097738), rabbit anti-cFos 1:500 (Synaptic Systems, 226008, RRID:AB_2737414), goat anti-SPO 1:5000 (Santa Cruz Biotechnology, 51212, RRID:AB_1129844), mouse anti-Kirrel3 1:1000 (Neuromab, 75-333, RRID:AB_2315857), mouse anti-GAPDH 1:2000 (Sigma Aldrich Millipore, MAB374, RRID:AB_2107445), rabbit anti-HA 1:1000 (Cell Signaling, 3724S, RRID:AB_1549585), and mouse anti-FLAG 1:1000 (Sigma Aldrich Millipore, F1804, RRID:AB_262044). All secondary antibodies were from Jackson ImmunoResearch Laboratories, made in donkey or goat, and used at 1:1000 for IHC and 1:2000 for Western blot.

### Viruses

To label mossy fiber boutons and filopodia, we replaced the GFP from the plasmid pCAG-GFP (RRID:Addgene 16664) with a palmitoylated GFP for membrane targeting. The virus was packaged at the Drug Discovery Core of the University of Utah. AAV plasmids encoding pAAV-syn1-EGFP-2A-3xHA-Kirrel3 (WT, R161H, or R205Q) were designed by the Williams lab and then constructed and packaged by VectorBuilder using the Kirrel3K isoform^46^. For synapse sparse labeling, AAVs were used at 10^9^ GC/mL and for Kirrel3 overexpression, AAVs were used at 10^13^ GC/mL. All AAVs were serotype PHP.eB.

### Western blot

Proteins were run on 8% acrylamide gels and transferred to nitrocellulose membranes using the Bio-Rad Turbo Transfer system. Membranes were incubated in blocking solution (10 mM Tris pH 7.5, 300 mM NaCl, 4% dry milk powder, and 0.1% Tween-20) for 10–15 min on a shaker, followed by primary antibody overnight at 4°C. They were washed three times in blocking solution, incubated in HRP-conjugated secondary antibodies (Jackson ImmunoResearch Laboratories) for 1-2 hours at room temperature, and detected using the Bio-Rad Clarity ECL kit (Cat#1705061) on a Bio-Rad ChemiDoc imaging system (RRID:SCR_019690). To prepare hippocampal lysates, 100 mg of hippocampal tissue was homogenized in 1 ml of 1x reducing sample buffer prepared from 4x stock consisting of 200mM Tris pH 6.8, 400mM DTT, 8% SDS, 40% glycerol, and 0.4% bromophenol blue.

### Fluorescent in situ hybridization

Fluorescent in situ hybridization chain reaction (Molecular Instruments HCR v3.0) was performed as previously described^47^. Briefly, 30 μm mouse brain cryo-sections were fixed in 4% PFA and washed in 1xPBS. Slices were treated with 0.05 mg/mL proteinase K then washed in 2x SSC buffer. HCR probes were designed and generated by Molecular Instruments for Gad1 and Kirrel3. After nuclear staining with Hoechst in 5x SSC-Tween-20, coverslips were mounted in Fluoromount-G and imaged on a confocal microscope. All reagents were prepared using 1x DEPC treated water.

### Foot shock-evoked cFOS Assay

Mice were subjected to a foot shock protocol consisting of three shocks delivered 1 minute apart. After the final shock, mice remained in the chamber for 30 min after shock onset, when brains were harvested. The foot shock current intensity was 0.5 mA for P14 and 1.0 mA for P30 and adult mice, with chamber lights on and no additional sensory stimuli. Shock delivery was verified by visual observation of the mouse reactions from video recordings. Home cage mice were not placed in the shock chamber. Perfusion and immunohistochemistry were performed as above. Imaging was conducted on an LSM980 confocal microscope (RRID:SCR_025048) using a 1 µm Z-stack interval, acquiring 15–20 optical sections per z-stack. ImageJ (RRID:SCR_002285) was used to quantify the density of cFOS-positive CA3 neurons per CA3 area (mm^2^). Maximum intensity z-projections were analyzed. CA3 area was calculated by drawing the boundaries of CA3 pyramidal layer using the Hoechst channel. For cFOS quantification, cFOS channel was thresholded manually by an experimenter blind to condition and genotype. The “analyze particles” command was used to count the number of cFOS particles in the CA3 cell body layer.

### MFB and filopodia synapse analysis

Neonatal pups (P0-2) underwent intra-cerebroventricular injections of AAV as described^48^. At P14, mice were perfused, brains sectioned, and immunohistochemistry was done as described above. Mossy fiber synapses (boutons and their filopodia) from the supra-pyramidal bundle were imaged on an LSM980 Airy scan microscope using the 63x objective, 5x zoom, and a 0.15 µm Z-stack interval through the entire synapse. Finally, Zeiss Zen Blue v3.7 software (RRID:SCR_013672) was used to generate 3D Airy scan reconstructions of mossy fiber boutons and filopodia for quantification of filopodia synapse density (marked by synaptoporin signal). Filopodia synapses were determined by analyzing the number of synaptoporin puncta within GFP-labeled filipodia.

### Neuron culture

Neuron cultures were prepared from P0 rat hippocampus. 18 mm borosilicate glass coverslips (Fisher, catalog #12-541-006 or Carolina, catalog #633033) were baked at approximately 200C for 5 hours then coated in 0.05 mg/mL poly-D-Lysine (Millipore Sigma, catalog #A-003-M) diluted in 0.1M Borate Buffer, pH 8.5. Coverslips were washed three times in autoclaved ddH₂O and equilibrated in plating medium before use. P0 rat hippocampi (both sexes) were dissected in ice-cold HBSS (10X HBSS Gibco #14185-052) supplemented with 2.5 mM HEPES, pH 7.4, 30 mM glucose, 1 mM CaCl2,1mM MgSO4, and 4 mM NaHCO3. Hippocampi were dissociated by incubating in papain solution (20 units/ml papain (Worthington, catalog #3126), 82 mM Na2SO4,30 mM K2SO4,5.8 mM MgCl2, 0.25 mM CaCl2,1mM HEPES, pH7.4, 20 mM glucose, and 0.1 mM NaOH) for 30 min at 37°C followed by trituration through a polished glass pipette. Dissociated cells were diluted in neuron plating media: MEM supplemented with 10% horse serum (Invitrogen, catalog #26050-088), 20 mM glucose, 100 units penicillin-streptomycin solution (PenStrep, Gibco, catalog #15140-122), 2 mM Glutamax (Invitrogen, catalog #35050-061), and 1 mM sodium pyruvate. Neurons were plated at 1.5 x 10^5^ cells/well onto glass coverslips in a 12-well plate. Neurons were cultured at 37°C and 5% CO2. 1-2 hours after plating, neuron plating media was replaced with feeding media: Neurobasal-A Plus (Gibco, catalog #A3582901) supplemented with B-27 Plus Supplement (50X) (Gibco, catalog #A3582801), 25 units penicillin-streptomycin solution, 0.5 mM Glutamax, and 12 mM glucose. Neurons were transfected on day 4 using the calcium phosphate method.

### Experimental design and statistical analysis

Statistics were calculated in GraphPad Prism (<u>graphpad.com</u>, RRID:SCR_002798). A priori power analysis was performed using G*Power 3.1 software (RRID:SCR_013726)^49^ using preliminary data and prior work to estimate sample sizes. Samples were analyzed blind to genotype or treatment. Sample sizes and statistical tests are reported in the figure legends and a statistics table (Supplementary Table 2). For nested analyses, mouse was the biological replicate, and mossy fiber boutons or sections were technical observations nested within each mouse. Dunnett’s pairwise multiple comparisons were done using the WT/control condition for comparison.

## DATA AVAILABILITY STATEMENT

The CA3 cFOS datasets generated and analyzed during this study are deposited in Hive and are publicly available at https://doi.org/10.7278/S5d-sw4s-gey8. Source data for all other data sets generated in this study are available from the corresponding author.

## ACKNOWLEDGEMENTS

We thank all members of the Williams lab and Dr. Dimitri Tränkner for feedback and discussion. This work was supported by a University of Utah Graduate Research Fellowship (O.S.) and NIH grants R01MH125943 (M.W.) and RF1MH134515 (M.W.).

---

**Supplement to Figure 1.**
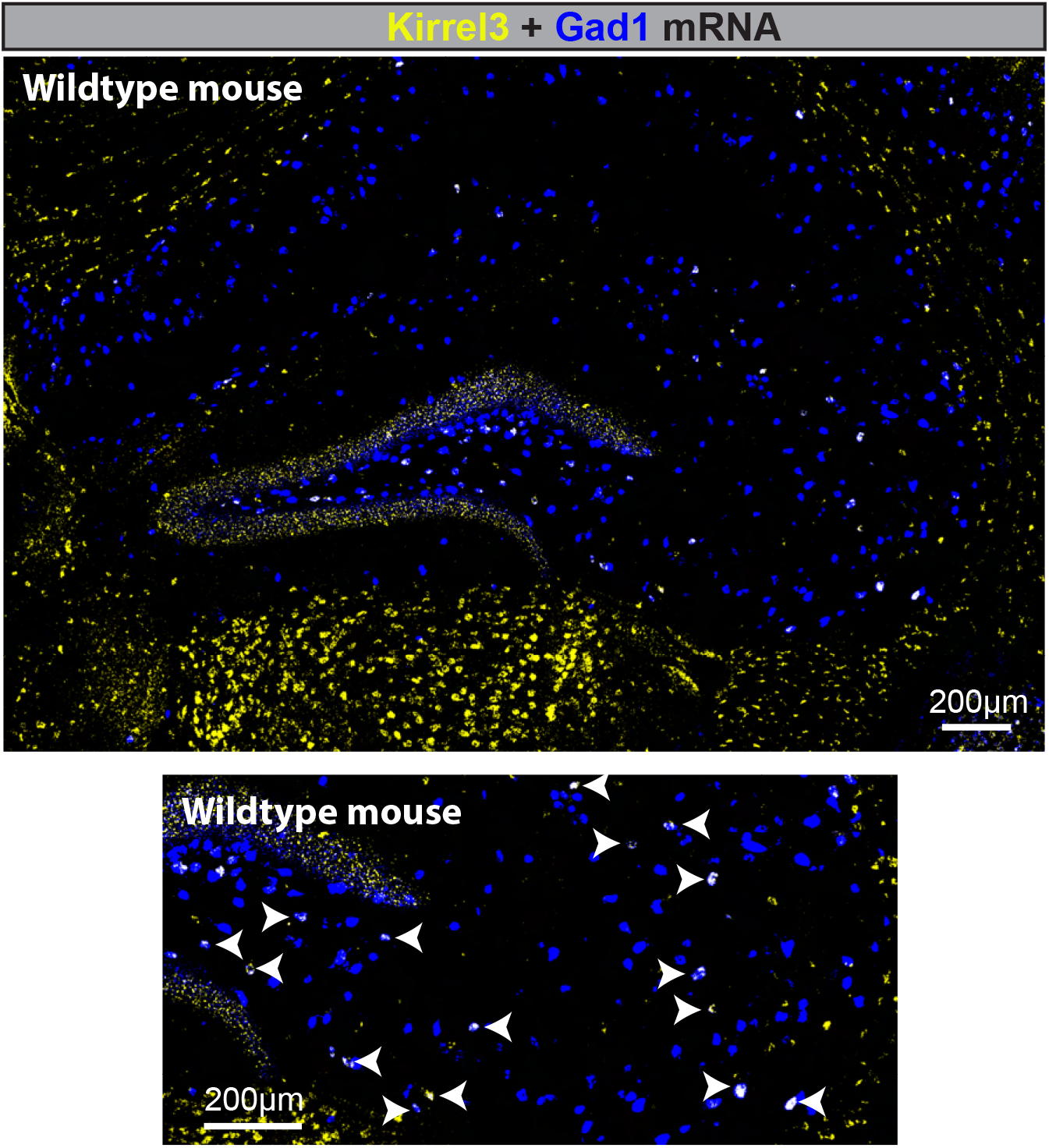
Kirrel3 mRNA expression in wildtype hippocampus. Wildtype hippocampal section stained via fluorescence in situ hybridization for mRNA of Kirrel3 (yellow) and the GABA neuron marker Gad1 (blue). Bottom image is a magnified image showing scattered Kirrel3 expressing-GABA neurons indicated by the arrow heads.

**Supplement to Figure 3.**
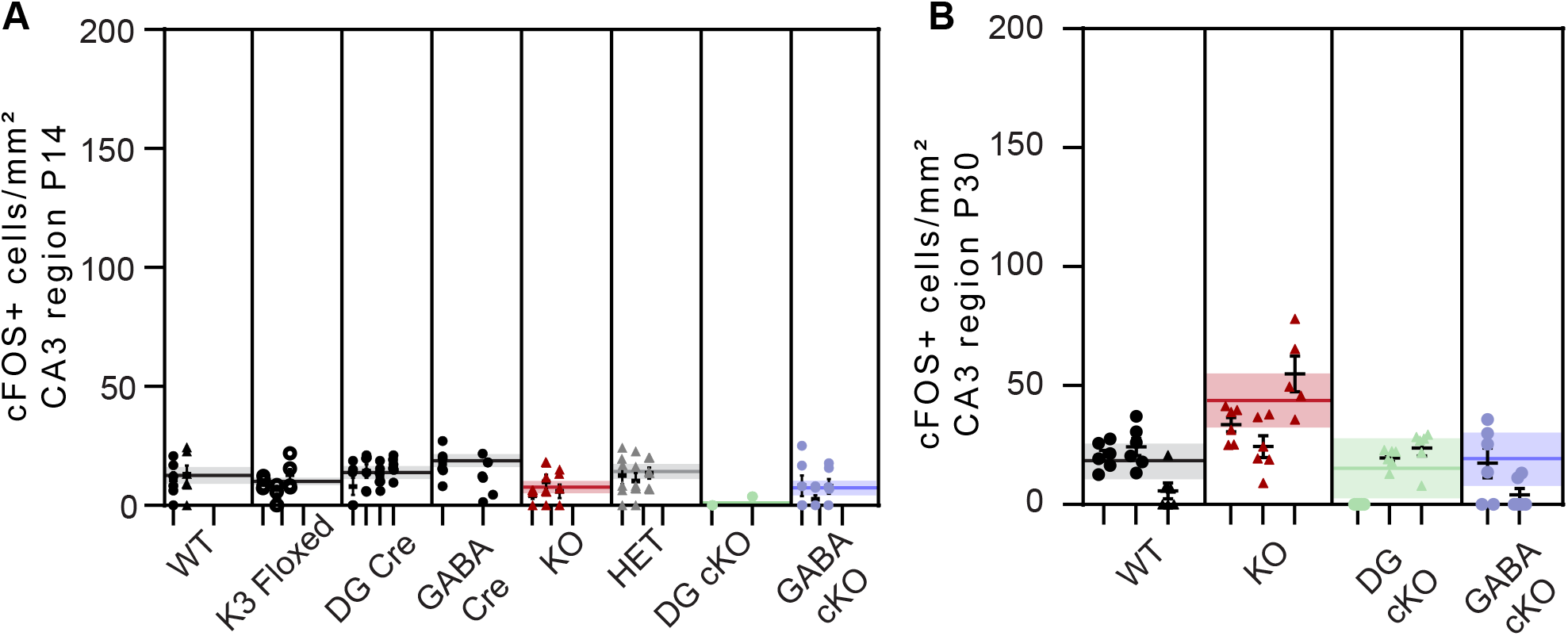
CA3 cFOS quantification in the home cage condition. **A-B)** Quantification of the density of cFOS+ CA3 neurons in mice from each genotype that were perfused directly from their home cage (unstimulated) at P14 and P30. Each mouse is shown in a column with 6 brain sections (dots) per mouse. Males are represented by a triangle and females a circle. The horizontal line and shading across each genotype is the global average and SEM. Nested one-way ANOVA indicates a significant difference in A (p=0.027) but Dunnett’s pairwise multiple comparisons indicate no significant differences for any genotype compared to WT control mice. ANOVA indicates no significant differences for B.

**Supplement to Figure 5.**
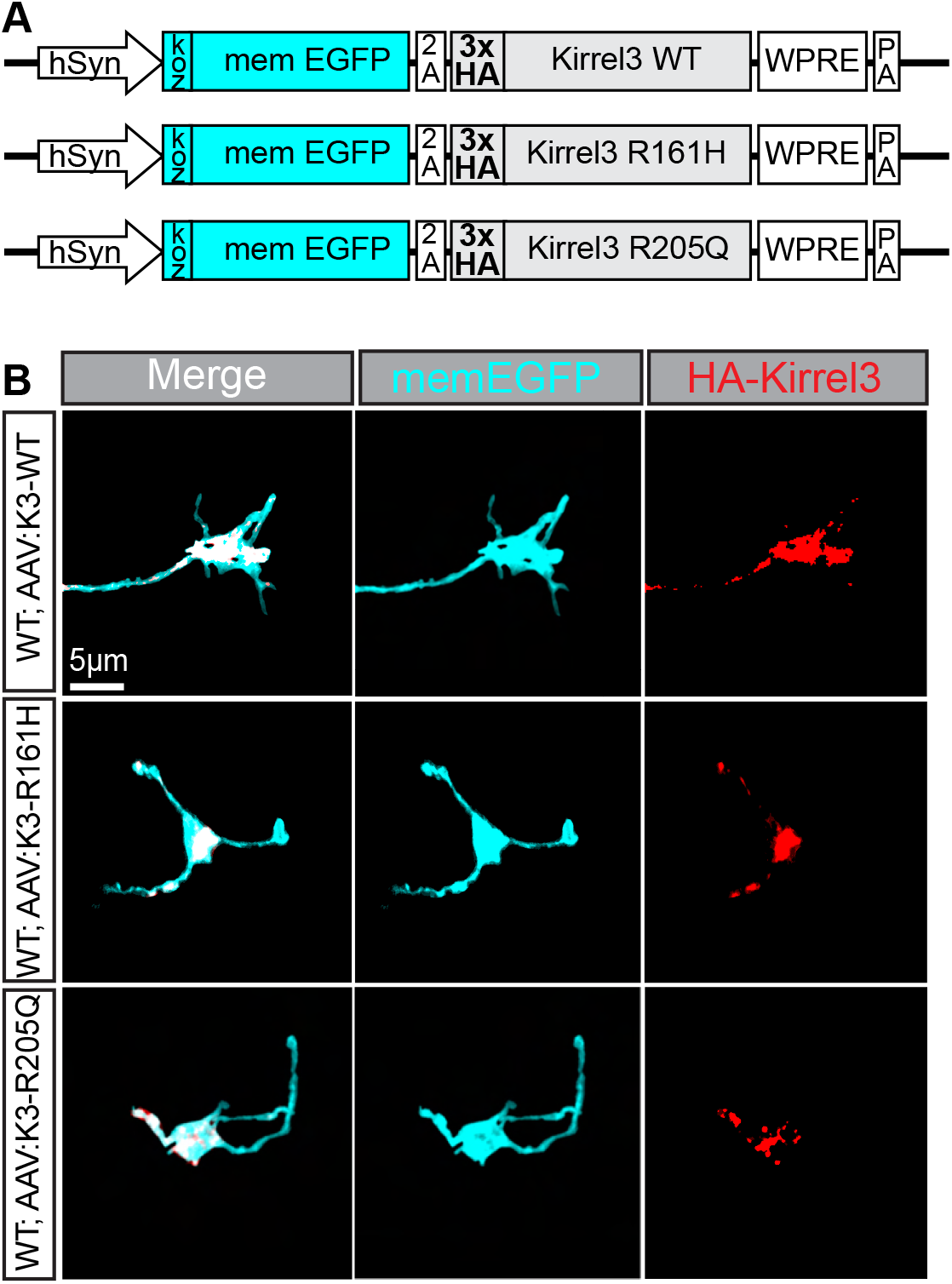
Viral-mediated Kirrel3 expression in DG mossy fiber axons. **A)** Diagram of AAV constructs used to overexpress Kirrel3 wildtype or R161H and R205Q variants. All were made using the Kirrel3K isoform. **B)** Representative images of immunostained MFBs show the membrane GFP (cyan) and HA-tagged Kirrel3 (red) in the MFB and filopodia.

**Supplementary Table 1:**
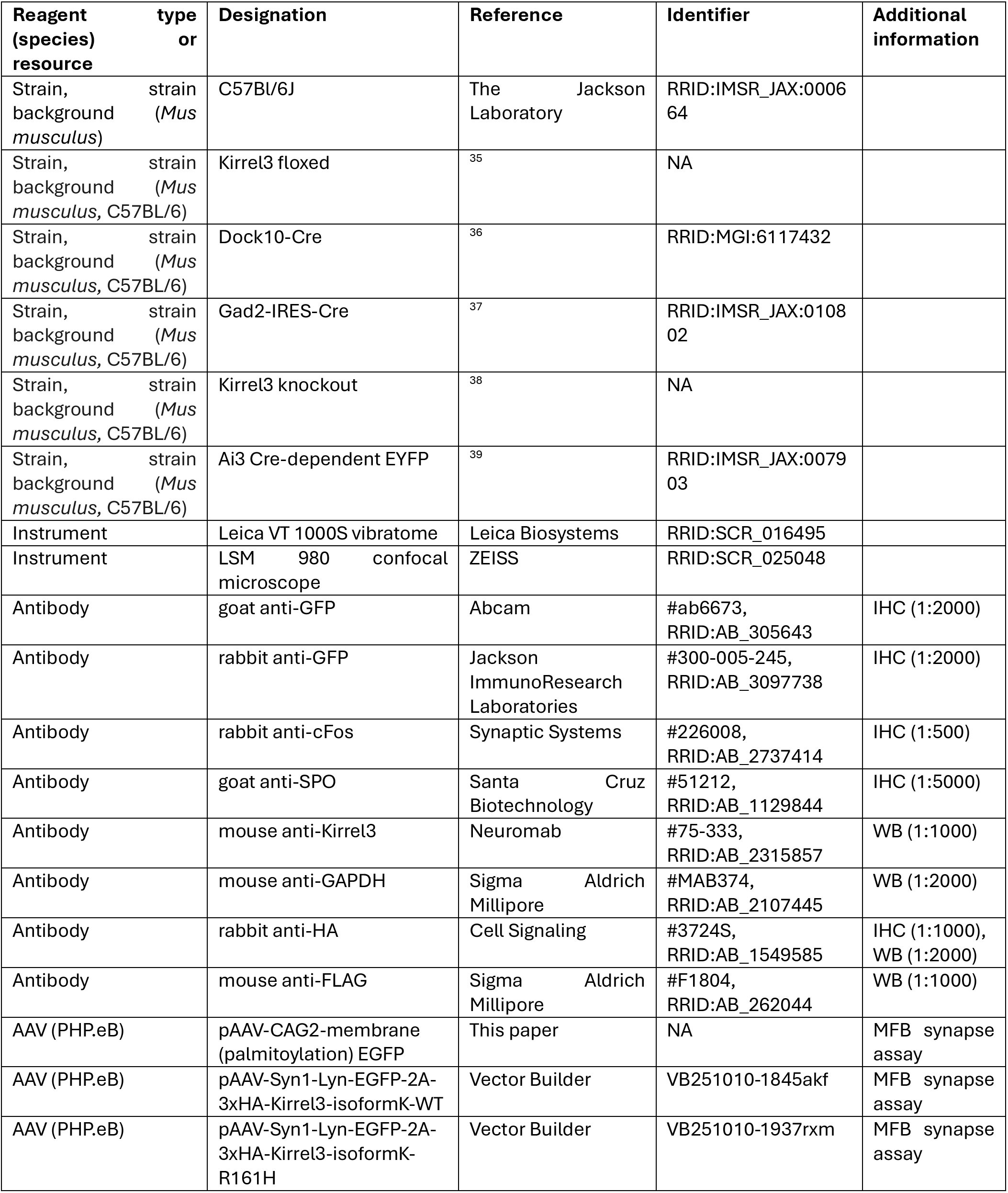
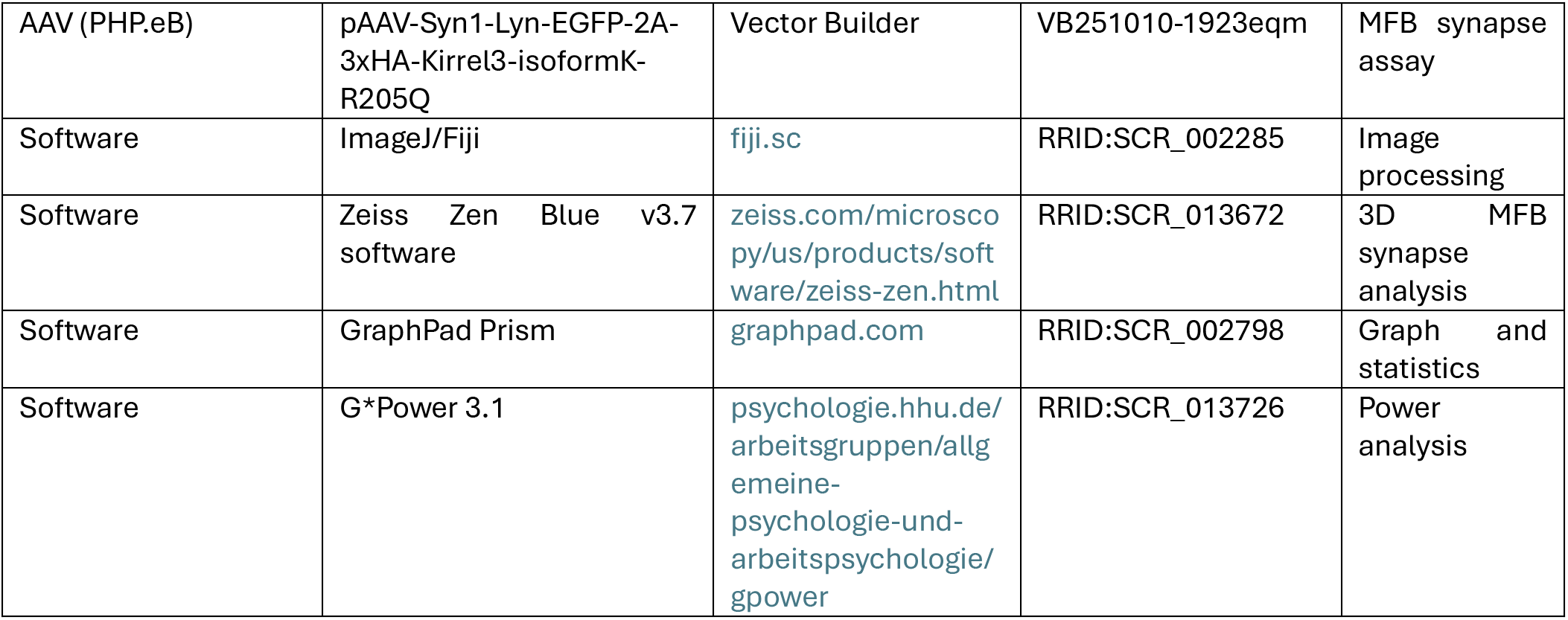
key resources.

**Supplementary Table 2:** Statistics.

| Figure | Statistical test used | Group labels (G1-Gn) | #Treatments | #Subjects | #Values | overall nested ANOVA p value | F, DFn (degrees freedom) numerator, Dfd | Significance |
| --- | --- | --- | --- | --- | --- | --- | --- | --- |
| 2D | Nested one-way ANOVA | Floxed, DG cKO, GABA, cKO | NA | 3,3,5 | 152 | 0.0025 | 14, 2, 8 | yes |
| 3C | Nested one-way ANOVA | Floxed, DG Cre, GABA Cre, DG cKO, GABA, cKO | NA | 6,6,6,6,6 | 180 | <0.0001 | 37, 4, 25 | yes |
| 3G | Nested one-way ANOVA | WT, KO, DG cKO, GABA cKO | NA | 6,6,6,6 | 138 | 0.0056 | 5.7, 3, 20 | yes |
| 4B | Nested one-way ANOVA | WT, KO, HET | NA | 4,5,4 | 156 | 0.0046 | 9.7, 2, 10 | yes |
| 4E | Nested one-way ANOVA | WT, KO, HET | NA | 6,6,6 | 107 | <0.0001 | 90, 2, 15 | yes |
| 6B | Nested one-way ANOVA | GFP, WT, R161H, R205Q | NA | 3,4,6,6 | 188 | 0.0016 | 8.5, 3, 15 | yes |
| 6E | Nested one-way ANOVA | GFP, WT, R161H, R205Q | NA | 5,6,6,6 | 137 | <0.0001 | 15, 3, 19 | yes |
| S3A | Nested one-way ANOVA | WT, Floxed, DG Cre, GABA Cre, KO, HET, DG cKO, GABA cKO | NA | 2,3,4,2,3,3,2,3 | 132 | 0.027 | 3.3, 7, 14 | yes |
| S3B | Nested one-way ANOVA | WT, KO, DG cKO, GABA cKO | NA | 3,3,3,2 | 65 | 0.13 | 2.6, 3, 7 | no |

## Notes

### Competing Interest Statement

The authors have declared no competing interest.

## REFERENCES

1. Südhof, T.C. Towards an Understanding of Synapse Formation. Neuron 100, 276–293 (2018).

2. Dalva, M.B., McClelland, A.C. & Kayser, M.S. Cell adhesion molecules: signalling functions at the synapse. Nat Rev Neurosci 8, 206–220 (2007).

3. Südhof, T.C. The cell biology of synapse formation. J Cell Biol 220(2021).

4. Missler, M., Südhof, T.C. & Biederer, T. Synaptic cell adhesion. Cold Spring Harb Perspect Biol 4, a005694 (2012).

5. Yogev, S. & Shen, K. Cellular and molecular mechanisms of synaptic specificity. Annu Rev Cell Dev Biol 30, 417–437 (2014).

6. Sperry, R.W. CHEMOAFFINITY IN THE ORDERLY GROWTH OF NERVE FIBER PATTERNS AND CONNECTIONS. Proc Natl Acad Sci U S A 50, 703–710 (1963).

7. Moreland, T. & Poulain, F.E. To Stick or Not to Stick: The Multiple Roles of Cell Adhesion Molecules in Neural Circuit Assembly. Frontiers in Neuroscience **Volume** 16 **-** 2022(2022).

8. Lepeta, K., et al. Synaptopathies: synaptic dysfunction in neurological disorders - A review from students to students. J Neurochem 138, 785–805 (2016).

9. van Spronsen, M. & Hoogenraad, C.C. Synapse pathology in psychiatric and neurologic disease. Curr Neurol Neurosci Rep 10, 207–214 (2010).

10. Shennib, O., Raines, O., Karamian, A.S. & Williams, M.E. Evaluation of the synapse adhesion molecule Kirrel3 in neurological disease. Front Neurol 16, 1662931 (2025).

11. Qi, C., Luo, L.D., Feng, I. & Ma, S. Molecular mechanisms of synaptogenesis. Front Synaptic Neurosci 14, 939793 (2022).

12. Gonçalves-Martins, J., Cação, C. & Ferreira, J.S. Neurexins, Ephrin receptors, and N-cadherin signaling as emerging mechanisms in synaptic dysfunction and neurodegenerative diseases. Commun Biol **9**(2026).

13. Shen, K. & Bargmann, C.I. The immunoglobulin superfamily protein SYG-1 determines the location of specific synapses in C. elegans. Cell 112, 619–630 (2003).

14. Sugie, A., Umetsu, D., Yasugi, T., Fischbach, K.F. & Tabata, T. Recognition of pre- and postsynaptic neurons via nephrin/NEPH1 homologs is a basis for the formation of the Drosophila retinotopic map. Development 137, 3303–3313 (2010).

15. Helmstädter, M., Höhne, M. & Huber, T.B. A brief overview on IRM function across evolution. J Neurogenet 28, 264–269 (2014).

16. Gerke, P., et al. Neuronal expression and interaction with the synaptic protein CASK suggest a role for Neph1 and Neph2 in synaptogenesis. J Comp Neurol 498, 466–475 (2006).

17. Wanner, N., et al. Functional and spatial analysis of C. elegans SYG-1 and SYG-2, orthologs of the Neph/nephrin cell adhesion module directing selective synaptogenesis. PLoS One 6, e23598 (2011).

18. Shen, K., Fetter, R.D. & Bargmann, C.I. Synaptic specificity is generated by the synaptic guidepost protein SYG-2 and its receptor, SYG-1. Cell 116, 869–881 (2004).

19. Özkan, E., et al. Extracellular architecture of the SYG-1/SYG-2 adhesion complex instructs synaptogenesis. Cell 156, 482–494 (2014).

20. Martin, E.A., et al. The intellectual disability gene Kirrel3 regulates target-specific mossy fiber synapse development in the hippocampus. Elife 4, e09395 (2015).

21. Serizawa, S., et al. A neuronal identity code for the odorant receptor-specific and activity-dependent axon sorting. Cell 127, 1057–1069 (2006).

22. Taylor, M.R., et al. Kirrel3-Mediated Synapse Formation Is Attenuated by Disease-Associated Missense Variants. J Neurosci 40, 5376–5388 (2020).

23. Wang, J., et al. Molecular and structural basis of olfactory sensory neuron axon coalescence by Kirrel receptors. Cell Rep 37, 109940 (2021).

24. Martin, E.A., Woodruff, D., Rawson, R.L. & Williams, M.E. Examining Hippocampal Mossy Fiber Synapses by 3D Electron Microscopy in Wildtype and Kirrel3 Knockout Mice. eNeuro 4(2017).

25. Hisaoka, T., Komori, T., Fujimoto, K., Kitamura, T. & Morikawa, Y. Comprehensive expression pattern of kin of irregular chiasm-like 3 in the adult mouse brain. Biochem Biophys Res Commun 563, 66–72 (2021).

26. Guo, N., et al. Dentate granule cell recruitment of feedforward inhibition governs engram maintenance and remote memory generalization. Nat Med 24, 438–449 (2018).

27. Lawrence, J.J. & McBain, C.J. Interneuron diversity series: containing the detonation--feedforward inhibition in the CA3 hippocampus. Trends Neurosci 26, 631–640 (2003).

28. Tuñon-Ortiz, A., et al. Inhibitory Neurons Marked by the Connectivity Molecule Kirrel3 Regulate Memory Precision. J Neurosci 45(2025).

29. Querzani, A., et al. KIRREL3-related disorders: a case report confirming the radiological features and expanding the clinical spectrum to a less severe phenotype. Ital J Pediatr 49, 99 (2023).

30. Guerin, A., et al. Interstitial deletion of 11q-implicating the KIRREL3 gene in the neurocognitive delay associated with Jacobsen syndrome. Am J Med Genet A 158a, 2551–2556 (2012).

31. Bhalla, K., et al. Alterations in CDH15 and KIRREL3 in patients with mild to severe intellectual disability. Am J Hum Genet 83, 703–713 (2008).

32. Ciaccio, C., et al. A Missense De Novo Variant in the CASK-interactor KIRREL3 Gene Leading to Neurodevelopmental Disorder with Mild Cerebellar Hypoplasia. Neuropediatrics 52, 484–488 (2021).

33. De Rubeis, S., et al. Synaptic, transcriptional and chromatin genes disrupted in autism. Nature 515, 209–215 (2014).

34. Narayan, P.R., et al. Identifying the Potential Role of Missense KIRREL3 Variants in Neurodevelopmental Phenotypes: A Case Series. Am J Med Genet A (2026).

35. Völker, L.A., et al. Neph2/Kirrel3 regulates sensory input, motor coordination, and home-cage activity in rodents. Genes Brain Behav 17, e12516 (2018).

36. Kohara, K., et al. Cell type-specific genetic and optogenetic tools reveal hippocampal CA2 circuits. Nat Neurosci 17, 269–279 (2014).

37. Taniguchi, H., et al. A resource of Cre driver lines for genetic targeting of GABAergic neurons in cerebral cortex. Neuron 71, 995–1013 (2011).

38. Prince, J.E., Brignall, A.C., Cutforth, T., Shen, K. & Cloutier, J.F. Kirrel3 is required for the coalescence of vomeronasal sensory neuron axons into glomeruli and for male-male aggression. Development 140, 2398–2408 (2013).

39. Madisen, L., et al. A robust and high-throughput Cre reporting and characterization system for the whole mouse brain. Nat Neurosci 13, 133–140 (2010).

40. Williams, M.E., et al. Cadherin-9 regulates synapse-specific differentiation in the developing hippocampus. Neuron 71, 640–655 (2011).

41. Singec, I., et al. Synaptic vesicle protein synaptoporin is differently expressed by subpopulations of mouse hippocampal neurons. J Comp Neurol 452, 139–153 (2002).

42. Torborg, C.L., Nakashiba, T., Tonegawa, S. & McBain, C.J. Control of CA3 output by feedforward inhibition despite developmental changes in the excitation-inhibition balance. J Neurosci 30, 15628–15637 (2010).

43. Hoffman, G.E., Smith, M.S. & Verbalis, J.G. c-Fos and related immediate early gene products as markers of activity in neuroendocrine systems. Front Neuroendocrinol 14, 173–213 (1993).

44. Bonapersona, V., et al. The mouse brain after foot shock in four dimensions: Temporal dynamics at a single-cell resolution. Proc Natl Acad Sci U S A 119(2022).

45. Lin, X., et al. c-Fos mapping of brain regions activated by multi-modal and electric foot shock stress. Neurobiol Stress 8, 92–102 (2018).

46. Traenkner, D., et al. Modular Splicing Is Linked to Evolution in the Synapse-Specificity Molecule Kirrel3. eNeuro 10(2023).

47. Choi, H.M.T., et al. Third-generation in situ hybridization chain reaction: multiplexed, quantitative, sensitive, versatile, robust. Development 145(2018).

48. Kim, J.Y., Grunke, S.D., Levites, Y., Golde, T.E. & Jankowsky, J.L. Intracerebroventricular viral injection of the neonatal mouse brain for persistent and widespread neuronal transduction. J Vis Exp, 51863 (2014).

49. Faul, F., Erdfelder, E., Lang, A.G. & Buchner, A. G*Power 3: a flexible statistical power analysis program for the social, behavioral, and biomedical sciences. Behav Res Methods 39, 175–191 (2007).

